# Detection of alternative splicing: deep sequencing or deep learning?

**DOI:** 10.1101/2025.08.23.671909

**Authors:** Lena Maria Hackl, Fabian Neuhaus, Sabine Ameling, Uwe Völker, Jan Baumbach, Olga Tsoy

**Affiliations:** Institute for Computational Systems Biology, University of Hamburg, Albert-Einstein-Ring 8-10, 22761 Hamburg, Germany; Interfaculty Institute for Genetics and Functional Genomics, University Medicine Greifswald, Felix-Hausdorff-Straße 8, 17475 Greifswald, Germany; Institute of Mathematics and Computer Science, University of Southern Denmark, Campusvej 55, 5230 Odense, Denmark; Department of Computer Science, Bioinformatics, Vrije Universiteit Amsterdam, De Boelelaan 1105, 1081 HV, Amsterdam, The Netherlands

## Abstract

Alternative splicing (AS) is a central mechanism of gene regulation that enables condition- and tissue-specific expression of gene isoforms. Its dysregulation plays a role in diseases such as cancer, neurological disorders, and metabolic conditions. Despite its importance, accurately identifying AS events remains challenging, especially in large-scale studies relying on publicly available RNA sequencing (RNA-seq) data. State-of-the-art AS event detection typically requires deep sequencing with over 100 million reads; however, much of the publicly accessible data is of lower sequencing depth. Recent advances, particularly deep learning models working with genomic sequences, offer new avenues for predicting AS without reliance on high sequencing depth data. Our study addresses the question: Can we utilize the vast repository of publicly available RNA-seq data for AS detection, despite often lacking the sequencing depth typically required? We show that sequence-based tools such as DeepSplice and SpliceAI show promising performance in retrieving novel and unannotated splice junctions, even when RNA-seq data are limited, but are not suitable for *de novo* splice junction detection. Our results demonstrate the potential of sequence-based tools for initial hypothesis development and as additional filters in standard RNA-seq pipelines, especially when sequencing depth is limited. Nonetheless, validation with higher sequencing depths remains essential for confirmation of splice events. Overall, our findings underscore the need for integrative methods combining genomic and RNA-seq data for prediction of tissue-/condition-specific AS in resource-limited settings.

## Introduction

### Cracking the splicing code for alternative splicing prediction

Alternative splicing expands the plasticity and functional diversity of the proteome, as a single gene can generate multiple protein isoforms tailored to cellular context. However, this plasticity requires fine-tuned regulatory mechanisms as aberrant splicing events can lead to the production of non-functional or even harmful protein isoforms, contributing to disease pathogenesis. Dysregulation of alternative splicing is associated with cancer [1–3] and, among others, neurological and metabolic diseases [4].

Alternative splicing is regulated by an intricate interplay of trans-acting proteins — splicing factors and spliceosome components — binding to the corresponding cis-acting pre-mRNA regulatory elements to either repress or activate the inclusion of nearby exons/introns [5]. Cis-acting elements include splice junctions, branch point sequences, and splicing enhancers and silencers. Additionally, epigenetic changes, such as histone modifications or chromatin structure, also influence splicing factor binding. The collection of rules that the cell employs to transform the information contained in a pre-mRNA sequence into a correctly spliced mRNA is referred to as the “splicing code” [6].

### Importance of RNA-seq sequencing depth for alternative splicing detection and prediction

RNA-sequencing (RNA-seq) is currently the most widely used method to measure the actual manifestation of alternative splicing events. However, different RNA-seq technologies have their advantages and limitations for alternative splicing detection. Short-read RNA sequencing technologies (Illumina and MGI [7]) allow for high-throughput detection and quantification of both known and novel splice variants, but do not capture full-length transcript isoforms. Long-read RNA sequencing technologies (Pacific Biosciences (PacBio) [8] and Oxford Nanopore Technologies (ONT) [9]) facilitate the detection of complex splicing events, but with a higher per-base error rate [10].

Regardless of the technology, sequencing depth is crucial for alternative splicing detection, especially of low-abundance transcripts. To quantify 80% of human transcripts with >10 FPKM, only around 36 million 100-bp paired-end reads are needed [11]. But, to detect rarer transcripts and alternative splicing events in human data, ∼100 to 150 million reads are necessary [12,13]. Moreover, detection of alternative splicing events from low sequencing depth data can be error-prone, with tools misidentifying or missing events for several reasons.

First, noise could be introduced during RNA library preparation. Sample degradation with time, contamination with non-target RNA, and variability in laboratory conditions can introduce misleading signals [14]. Sampling bias increases with low sequencing depth, and less common splice junctions might be covered by only a few or no reads due to a lower stochastic probability. Choice of the library construction protocol [15], experimental artifacts such as 3′–5′ transcript bias, random hexamer priming [16], or PCR amplification bias [17] can lead to under-representation of certain transcripts.

Second, noise could be introduced during sequencing. Isoforms that do not exist could be incorrectly called due to base-calling errors introducing wrong donor or acceptor sites. Furthermore, longer transcripts will be detected better than shorter ones as they generate more fragments and are better covered [18].

Third, noise could be introduced during subsequent computational processing, such as alignment of reads or alternative splicing event detection. Reads mapping to several positions lead to mapping ambiguity [19], especially in homologous gene families or repetitive regions. Therefore, the choice of a mapping and splice prediction tool introduces tool-specific biases.

Despite the difficulties of detecting splicing from low sequencing depth data, high sequencing depth data are only available to a limited extent. Most publicly accessible RNA-seq datasets, such as TCGA and GTEx, lack sufficient sequencing depth [13]. However, splicing prediction tools might open a possibility to utilize the extensive collection of these low sequencing depth data sets for comprehensive alternative splicing detection.

### Existing splice prediction tools

Splicing prediction tools can be categorized into tools predicting splice junctions [20], splice events [21,22], percent spliced in (PSI) value [23], aberrant splicing [24,25] or differential splicing [26].

In this work, we focus on tools that predict splice junctions. These splicing prediction tools can work with different types of input data.

First, there are tools that use DNA sequence data to predict splice junctions. The pioneers in this field used Hidden Markov Models [27] or Support Vector Machines [28]. Recently, deep learning architectures such as Convolutional Neural Networks [29], Long-Short Term Memory Neural Networks [30], and Recurrent Neural Networks [31], originally developed for natural language processing, have been applied to biological sequences to predict alternative splicing events. The most prominent examples are listed in Supplementary Table S1. They outperform conventional machine learning models in terms of automatically extracting sequence features and detecting non-linear correlations. By pretraining on big datasets, such models can learn and generalize knowledge for improved alternative splicing event prediction on limited, condition-specific datasets. Most commonly, the donor and acceptor site sequences with surrounding context are one-hot-encoded, and the neural network extracts sequence features that are then used for splicing prediction. Zhang et al. proposed the DeepSplice model based on a convolutional neural network that uses splice junctions and their surrounding sequence for training splice site prediction [20]. It was specifically developed to discern any novel splice junctions derived from RNA-seq alignment, but it does not take into account the number of reads covering a junction. A recent, already widely recognized, state-of-the-art tool, originally designed to predict the impact of genetic variants on splicing from sequence, is SpliceAI [29]. Part of its model architecture provides separate predictions for acceptor and donor sites, enabling it to directly predict splice junctions.

Second, there are tools that use RNA-seq data to predict splice junctions. Here, we distinguish between tools that merely quantify alternative splicing events from RNA-seq data and those that predict missing splicing events, and focus on the latter. To the best of our knowledge, only one tool, Junction Coverage Compatibility score (JCC), satisfies this criterion. The tool estimates the reliability of observed splice junction reads in RNA-seq data to identify spurious or novel junctions [32].

In our study, we focus on the specific question: Which of these two strategies is feasible for predicting alternative splicing events *de novo* and can overcome the necessity of high sequencing depth RNA-seq data? As the representative of the first category, we chose DeepSplice [20] as it was specifically designed to detect novel splice junctions from

RNA-seq alignment, and SpliceAI [29] due to its widespread adoption within the field. For the second category, we chose JCC [32] as it can find inconsistencies between RNA-seq data and expected coverage profiles to predict the reliability of a splice junction.

We investigated the feasibility of these tools to 1) detect splice junctions introduced by technical noise, 2) predict splice junctions that can not be detected from the low sequencing depth data, and 3) predict splice junctions that are neither annotated nor detected in low sequencing depth data.

The purpose of this study is not to benchmark existing splice prediction tools, as there are plenty of benchmarks already [33–35], but rather to explore the tools’ application to the aforementioned tasks. To our knowledge, this is the first study of its kind.

## Material and Methods

### Splice-aware RNA-seq aligners

A comparative study by Baruzzo et al. recommends using aligners Spliced Transcripts Alignment to a Reference (STAR), Hierarchical Indexing for Spliced Alignment of Transcripts (HISAT), HISAT2, or ContextMap2 for the detection of non-canonical junctions [36]. We decided on STAR (v2.7.10a) [19], which is commonly used in large-scale studies, and HISAT2 (v2.2.1) [37], as it shows even higher alignment sensitivity.

### Tools

First, we evaluated the feasibility of deep learning tools to extract splice junctions from DNA sequence. The most popular and recent tools here are SpliceAI [29] and DeepSplice [20], both based on convolutional neural networks (CNNs). SpliceAI (v1.3.1) [29], a 32-layer neural network based on the CNN ResNet architecture, uses a 10-kilobase flanking pre-mRNA sequence on each side of a position of interest as input to predict the probability of the position being a splice acceptor, donor, or neither. DeepSplice (v0.10) [20] models the donor and acceptor splice sites together as functional pairs to classify candidate splice junctions and filter out false-positive splice junctions.

Next, we evaluated the feasibility of tools to extract splice junctions from RNA-seq data and chose JCC (version August 2018) [32]. The JCC score reflects the reliability of transcript abundance estimates. By comparing the number of reads predicted to align across each junction (inferred from the transcript abundances and a fragment bias model) to the observed number of junction-spanning reads (obtained via alignment of the reads to the genome), it aims to find inconsistencies to differentiate between true-positive and false-positive splice junctions.

### Data

#### Decoy data set construction

As a set of positive splice junctions we used the combination of junctions from recount3 [38], and two RNA-seq data sets with approximately 500 million paired-end RNA-seq reads: 2 samples from adipose tissue [12] and 4 samples from patients with dilated cardiomyopathy (DCM).

Splice junctions are detected by short conserved sequence motifs, e.g., donor sites marked by GT and acceptor sites by AG. But throughout the genome, frequently similar non-functional sequences can be found. We created the decoy dataset by randomly sampling combinations of GT and AG positions from within the coding sequence and excluding those contained in the positive set. For this dataset of false-positive junctions, we preserved the distribution among the main chromosomes and the distribution of junction lengths from the positive set.

#### Gold standard data set construction

We aligned and extracted splice junctions from high sequencing depth RNA-seq data sets using STAR [19] and HISAT2 [37] (Figure 1a, step (1)) and kept only junctions on the main chromosomes. Next, we constructed four gold standard data sets based on different filtering criteria, as shown in Figure 1a, step (2). The first gold standard data set contains all junctions detected by STAR without any filtering. For the second data set, we filtered junctions detected by STAR according to Illumina guidelines [39]. The third data set is a result of the strictest filtering — filtering out all junctions detected by STAR with fewer than 10 uniquely aligned reads. The overlap between the different gold standards is shown in Supplementary Figure S1 and Supplementary Figure S2. Finally, the fourth data set contains all the junctions detected by HISAT2 without any filtering. We did not construct HISAT2 filtered gold standards, since the aligner does not provide the number of reads per junction.

**Figure 1:**
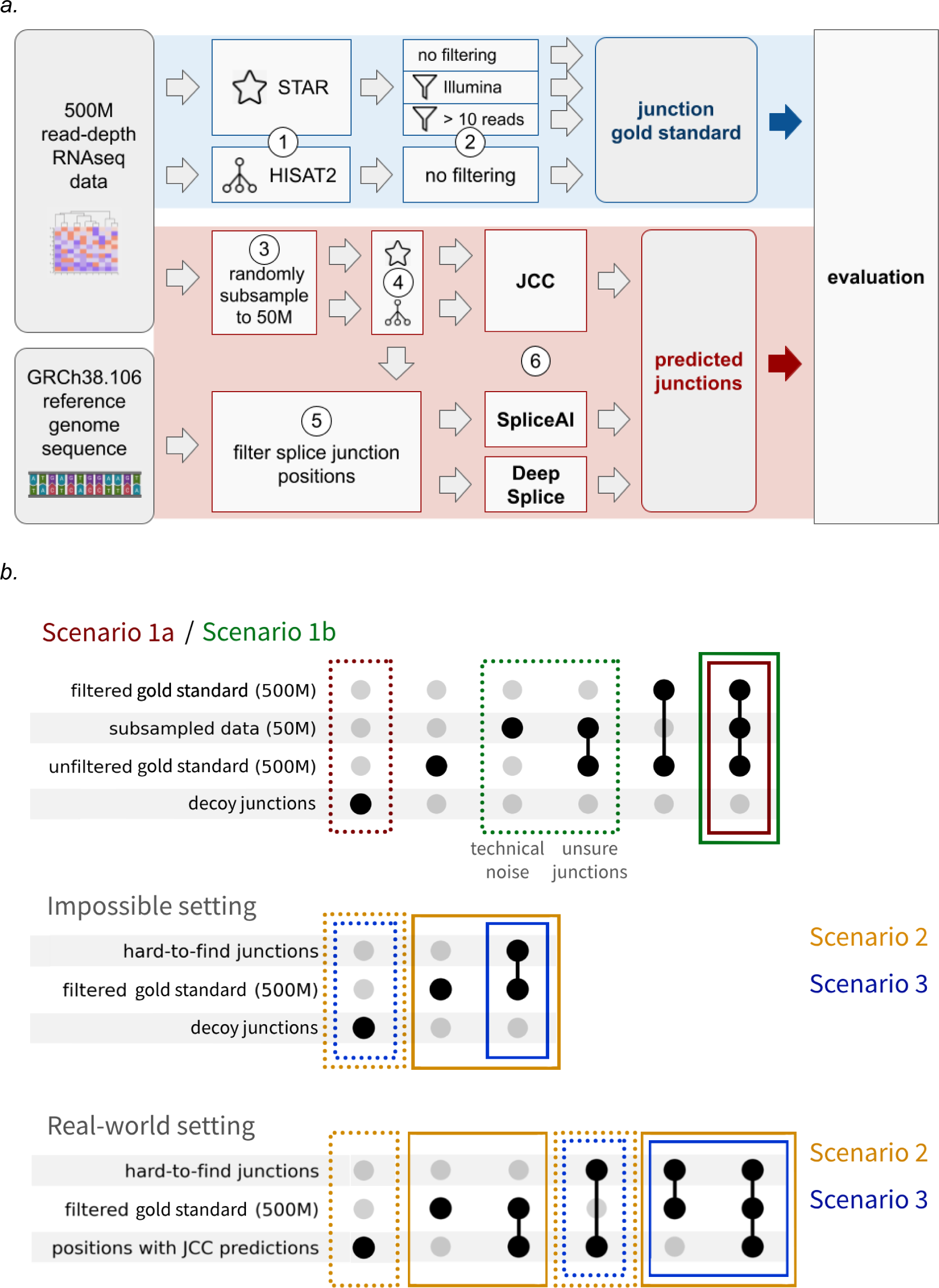
a. Analysis pipeline using the GRCh38 reference genome sequence and RNA-seq data with 500M reads from four DCM patients. (1) The reads are aligned using the two aligners STAR and HISAT2, (2) as STAR provides additional information the resulting junctions are filtered according to Illumina’s suggested criteria and filtered to only junctions with more than 10 uniquely mapping reads. (3) RNA-seq data is randomly subsampled ten times to samples of 50M reads, (4) the resulting reads are again aligned using the two aligners STAR and HISAT2. (5) According to these junction positions, surrounding sequences are extracted from the reference genome, and (6) tools SpliceAI, DeepSplice and JCC are run. Depending on the question we are answering and the corresponding evaluation scenario, we evaluate on a subset of all junctions the tools were run on, as described in panel b. The gold standard is marked in blue, and the tool predictions in red. b. The upset plot is showing the junction sets we evaluate the three tools on. The positive class is outlined with solid lines, whereas the negative class is outlined with dotted lines. For Scenario 1a: “Detecting junctions from a decoy dataset” (shown in red), we inverted the class labels, treating the decoy junctions as the positive class. We take junctions that are found in both the subsampled 50M reads data and the gold standard as negatives. For Scenario 1b: “Detecting spurious junctions” (shown in green), we inverted the class labels, treating the spurious junctions found only in the subsampled 50M reads data but not in the filtered gold standard as the positive class. We take junctions that are found in both the subsampled 50M reads data and the gold standard as negatives. For Scenario 2: “Predicting junctions that could be detected with higher sequencing depth” Hypothetical use case (shown in orange) and Scenario 3: “Predicting hard-to-find junctions” Hypothetical use case (shown in blue) as positives we take filtered gold standard positions, and as negatives decoy junctions. Here, hard-to-find junctions mean junctions from the filtered gold standard that were found neither in the subsampled 50M reads data nor in the reference genome. For Scenario 2: “Predicting junctions that could be detected with higher sequencing depth” Real-world use case (shown in orange) and Scenario 3: “Predicting hard-to-find junctions” Real-world use case (shown in blue), we take filtered gold standard positions and positions with JCC predictions. Here, hard-to-find junctions mean junctions from the filtered gold standard or predicted by JCC that were found neither in the subsampled 50M reads data nor in the reference genome.

#### Workflow

For input to the DNA sequence-based tools, we randomly subsampled the samples from the DCM patients 10 times to 50M reads. We aligned them to the human genome GRCh38, using the genome annotation from Ensembl 106, and called splice junctions using both HISAT2 and STAR as shown in Figure 1a, step (3). The overlap between the splice junctions detected by HISAT2 and STAR is shown in Supplementary Figure S3. For evaluation with SpliceAI and DeepSplice, we filtered the resulting junctions to a set of junction positions on the main chromosomes.

For input to SpliceAI, we extracted flanking sequences of 5,000 nucleotide length on either side of these junctions. We one-hot encoded and zero-padded the sequences, and calculated the SpliceAI acceptor and donor scores using the pre-trained model.

For input to DeepSplice, we extracted flanking sequences of 30 nucleotide length on either side of both donor and acceptor sites. We one-hot encoded those sequences, each consisting of four 30-nucleotide subsequences, and then ran the pre-trained DeepSplice model.

For input to JCC, we used STAR output, as HISAT2 output does not provide the number of reads per junction required to run JCC. JCC estimates the reliability of transcript-level abundance estimates per gene, which we transformed from a range of (2,0) to (0,1) and used as a reliability score for the splice junctions in this gene. JCC can be used with eight different transcript abundance estimation methods, but the results for each were found to be similar [32]. We combined GRCh38.106 cDNA and ncRNA references, built a transcriptome index, and estimated transcript abundance using Salmon (v1.10.0), R version 4.2.0 in quasi-mapping mode with sequence, GC, and positional bias correction. We then deduced inferential variance for each transcript using 100 bootstrap runs. To fit the bias model using alpine, we again used GRCh38.106 (Supplementary Text S2 for parameters).

While DeepSplice returns a single score per splice junction, JCC and SpliceAI return separate scores for donor and acceptor sites for each position. For evaluation, we combined these two scores by taking the maximum for each splice junction. More details can be found in Supplementary Text S1.

Note that the aligner used for the gold standard was always the same as the aligner used for the 50M input data mapping, and we did not perform a cross-comparison between the aligners.

For different evaluation scenarios, we selected subsets of junctions depending on the research question.

### Evaluation scenarios

In the study, we constructed evaluation scenarios (as shown in Figure 1 b) to address the following questions:

Scenario 1:

- a. What is the tools’ baseline performance for distinguishing true junctions from decoy junctions?
- b. Can the tools distinguish true splice junctions from spurious junctions?

Scenario 2:

- Can the tools predict splice junctions from 50M reads data that could only be found with higher sequencing depth?

Scenario 3:

- Can the tools predict splice junctions from 50M reads data that are not annotated in the reference genome and could only be found at higher sequencing depth?

To evaluate the tools’ performance, for each scenario we selected the corresponding subsets of splice junctions and calculated the area under the precision-recall curve (AUPRC) using average precision, as follows:

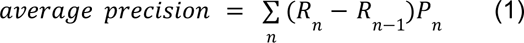

with 𝑃*_n_* and 𝑅*_n_* being precision and recall at the nth threshold:

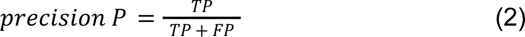

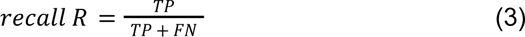

Given the pronounced class imbalance in several scenarios, we used the AUPRC metric, which is widely recommended for its focus on positive-class performance in imbalanced settings [40]. We scaled the DeepSplice scores to the range (0,1), to calculate F1 scores for differing thresholds n, as follows:

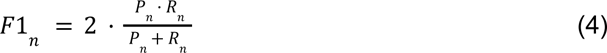

### Scenario 1: Detecting noisy junctions

#### a. Detecting junctions from a decoy dataset

Splice junctions are defined by consensus motifs, but these short canonical motifs (e.g., GT-AG) frequently occur in the genome by chance. Junctions based on

non-functional motifs constitute our decoy dataset. As a baseline, we evaluated the tools’ performance to distinguish well-supported splice junctions from decoy splice junctions. For this task, we inverted the class labels, treating the decoy junctions as the positive class.

We excluded JCC from this evaluation scenario because it requires read alignment information unavailable for junctions from the decoy dataset.

For SpliceAI and DeepSplice prediction, we used as input:

- sequences flanking splice junctions detected in both 50M and 500M reads data (negative class)
- the same number of sequences flanking junctions sampled from the decoy junctions (positive class).

#### b. Detecting spurious junctions

However, among sequences that resemble splice junctions, some even mislead mapping tools. Those sequences are detected as junctions at lower sequencing depth, while disappearing at higher sequencing depth. In this scenario, we evaluate the performance of the tools in detecting such spurious junctions introduced by technical noise (see

Supplementary Table S2) or by low-confidence signals that would typically be filtered out using cutoff or Illumina filtering criteria.

For this task, we again inverted the class labels, treating the spurious junctions as the positive class.

For JCC prediction, we used as input:

- splice junctions detected in both 50M and 500M reads data (negative class)
- splice junctions detected only in 50M but not in 500M reads data (positive class).

For SpliceAI and DeepSplice prediction, we used as input the sequences flanking those junctions.

#### Scenario 2: Predicting junctions that could be detected with higher sequencing depth

This scenario addresses the question whether we can use the underlying splicing code to retain information even from low sequencing depth RNA-seq data.

As JCC can redistribute RNA-seq reads to other positions on a gene, it can be run given only low sequencing depth RNA-seq data as input. This type of input corresponds to the input data most users might have. Thus, we named this part of the scenario “Real-world use case”. In contrast, DeepSplice and SpliceAI can not give predictions for positions that were not provided as input. Therefore, we provided all splice junctions of interest as input, including those that can only be detected from higher sequencing depth. Given that the user usually does not have this information, we named this part of the scenario “Hypothetical use case”.

For JCC prediction, we used as input:

- splice junctions detected in 50M reads data.

We evaluated JCC on all positions with predictions and on gold standard junctions. While we use gold standard annotations for evaluation, this use case represents the real-world situation where the user has no knowledge of splice junctions that could be detected from higher sequencing depth.

For SpliceAI and DeepSplice prediction, we used as input:

- sequences flanking splice junctions detected in 500M reads data (positive class)
- the same number of sequences flanking splice junctions from our decoy junction dataset (negative class).

We should note that this hypothetical use case does not represent a real-world situation. However, the design of SpliceAI and DeepSplice requires the pre-defined positions as input.

#### Scenario 3: Predicting hard-to-find junctions

This scenario addresses the most challenging case of retrieving novel hard-to-find splice junctions detected only in 500M but not in 50M reads data, and not annotated in the reference genome. These splice junctions might present rare and lowly expressed splicing events (see Supplementary Table S2).

For JCC prediction, we used as input:

- splice junctions detected in 50M reads data.

Here we evaluated JCC on hard-to-find splice junctions as positives and positions with JCC predictions but not in the input nor the gold standard as negatives.

As noted above, this part of the scenario is closer to the real-world use case.

For SpliceAI and DeepSplice prediction, we used as input:

- sequences flanking hard-to-find splice junctions (positive class)
- the same number of sequences flanking junctions from our decoy junction dataset (negative class).

This part of the scenario does represent the Hypothetical use case, rather than a real-world situation.

#### Evaluation metrics

In the results, we report the average and standard deviation of AUPRC for each scenario across 40 runs per tool (4 RNA-seq samples with 500M reads, each with 10 random subsamples with 50M reads). We used AUPRC, as several of our scenarios involve imbalanced datasets where precision-recall metrics better reflect positive class performance. For performance comparison, we implemented a baseline “No-Skill” classifier, which indiscriminately labels all inputs as positive. The AUPRC of this classifier is equal to the fraction of positives, and serves as the baseline for the other tools’ AUPRC. Supplementary Table S3 shows the number of junctions used for evaluating the scenarios described above.

## Results and Discussion

### Scenario 1: Detecting noisy junctions

#### a. Detecting junctions from a decoy dataset

We began with a baseline comparison examining the performance of tools in distinguishing between positions that have never been observed as splice junctions (see Section Decoy data set construction in Methods) and splice junctions detected in both 50M and 500M reads data (Table 1). We observed the best performance with AUPRCs of 0.99 for SpliceAI on the gold standard based on STAR output filtered with a specific cutoff (‘STAR cutoff’), but performance decreased strongly with less strict filtering. Overall, while both SpliceAI and DeepSplice surpassed the No-Skill baseline, SpliceAI consistently outperformed DeepSplice.

**Table 1:** The mean area under the precision and recall curve (AUPRC) values ± 1 standard deviation per tool are shown for Scenario 1a “Detecting junctions from a decoy dataset” and gold standards STAR without filtering, STAR with Illumina filtering, STAR with cutoff > 10 uniquely aligned reads and HISAT2 without filtering averaged across all 4 × 10 subsampled 50M reads data. “No-Skill” describes a baseline classifier that predicts all splice junctions as positives. To focus on the tools’ performance in identifying the negative class, we calculated AUPRC after inverting the class labels, treating false-positive junctions as the positive class.

|  | DeepSplice | SpliceAI | baseline classifier<br>("No-Skill") |
| --- | --- | --- | --- |
| **STAR unfiltered** | 0.0759 ± 4.3E-3 | 0.181 ± 0.012 | 0.001431 ± 8.3E-5 |
| **STAR Illumina** | 0.506 ± 0.025 | 0.711 ± 0.026 | 0.01499 ± 2.5E-4 |
| **STAR cutoff** | 0.97123 ± 4.0E-4 | 0.99512 ± 2.6E-4 | 0.2408 ± 2.9E-3 |
| **HISAT2 unfiltered** | 0.864 ± 0.016 | 0.9613 ± 5.5E-3 | 0.0427 ± 5.6E-3 |

The analysis demonstrated that, while the tools can differentiate decoy junctions from well-supported splice junctions, their performance decreases immensely for splice junctions with lower read support.

#### b. Detecting spurious junctions

In this evaluation scenario, we focus on spurious junctions. Those can be split up into 1) technical noise if they are found in the 50M but not in the 500M reads data, and 2) low confidence or low-expressed junctions if they are found in the 50M and 500M reads data but then filtered out based on any filtering criteria.

The technical noisy junctions introduced by misalignments of the mapping tools made up around 4.27% of all splice junctions detected by HISAT2 (on average 7.986 out of 186.920) and only around 0.14% of all splice junctions called by STAR (on average 359 out of 251.195). Both DeepSplice and SpliceAI show better performance than the No-Skill baseline classifier on all gold standard data sets (see Table 2). However, the No-Skill classifier outperforms JCC on the STAR Illumina gold standard. Similar to Scenario 1a, we observe improved tool performance with stricter filtering of the gold standard dataset. Generally, all tools are capable of distinguishing highly expressed junctions from spurious junctions, but perform poorly for the classification of lowly expressed junctions and spurious junctions.

**Table 2:**
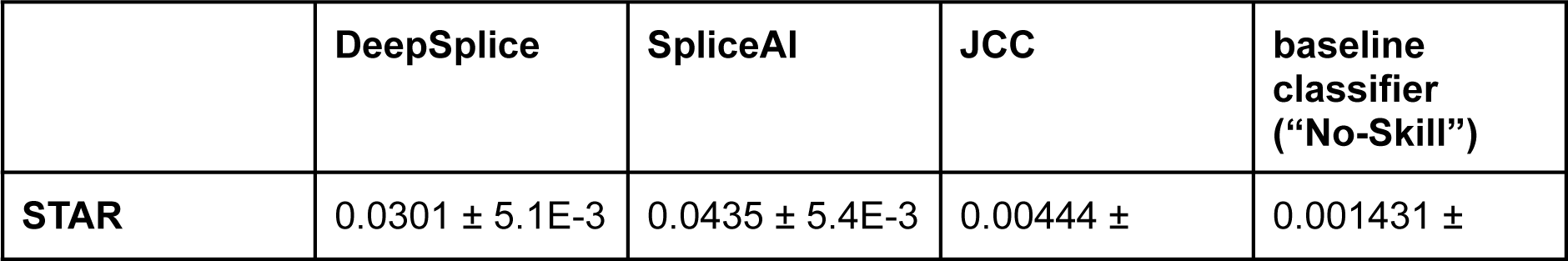
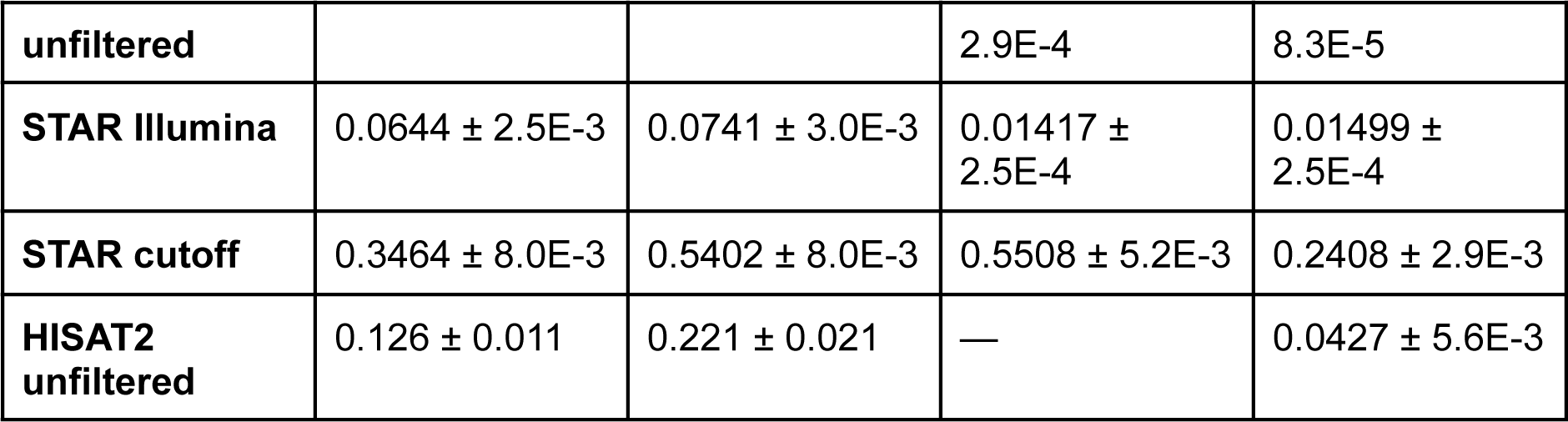
The mean area under the precision and recall curve (AUPRC) values ± 1 standard deviation per tool are shown for Scenario 1b “Detecting spurious junctions” and gold standards STAR without filtering, STAR with Illumina filtering, STAR with cutoff > 10 uniquely aligned reads and HISAT2 without filtering averaged across all 4 × 10 subsampled 50M reads data. “No-Skill” describes a baseline classifier that predicts all splice junctions as positives. To focus on the tools’ ability to detect noisy junctions, we calculated AUPRC after inverting the class labels, treating spurious junctions as the positive class.

### Scenario 2: Predicting junctions that could be detected with higher sequencing depth

In this scenario, we assessed the tools’ performance for predicting splice junctions that could be detected only at a higher sequencing depth. While JCC can re-distribute RNA-seq reads to the correct position using the actual data with lower sequencing depth (“Real-world use case”), DeepSplice and SpliceAI need all splice junctions of interest as input, which is usually not possible in real-world data analysis (“Hypothetical use case”).

While SpliceAI consistently outperforms DeepSplice with an AUPRC of 0.99 in comparison to 0.98 on the unfiltered gold standard STAR (Table 3, “Hypothetical use case”), the improvement is marginal. Performance improves with Illumina filtering, but worsens with even stricter cutoff filtering. Both tools outperform the No-Skill baseline classifier, which has an AUPRC of 0.5.

**Table 3:** The mean area under the precision and recall curve (AUPRC) values ± 1 standard deviation per tool are shown for Scenario 2 “Predicting junctions that could be detected with higher sequencing depth” and gold standards STAR without filtering, STAR with Illumina filtering, STAR with cutoff > 10 uniquely aligned reads and HISAT2 without filtering averaged across all 4 × 10 subsampled 50M reads data. “No-Skill” describes a baseline classifier that predicts all splice junctions as positives.

|  |  |  |
| --- | --- | --- |
|  | **JCC** | **baseline classifier (“No-Skill”)** |
| **STAR unfiltered** | 0.8508 ± 5.0E-3 | 0.8808 ± 4.6E-3 |
| **STAR Illumina** | 0.8268 ± 5.4E-3 | 0.8634 ± 5.0E-3 |
| **STAR cutoff** | 0.6622 ± 4.1E-3 | 0.6476 ± 2.2E-3 |

*Table 3: The mean area under the precision and recall curve (AUPRC) values ± 1 standard deviation per tool are shown for Scenario 2 "Predicting junctions that could be detected with higher sequencing depth" and gold standards STAR without filtering, STAR with Illumina filtering, STAR with cutoff > 10 uniquely aligned reads and HISAT2 without filtering averaged across all 4 × 10 subsampled 50M reads data.
|  | **DeepSplice** | **SpliceAI** | **baseline classifier**<br>**("No-Skill")** |
| --- | --- | --- | --- |
| **STAR unfiltered** | 0.9831 ± 1.7E-3 | 0.9927 ± 1.0E-3 | 0.5 ± 0.0 |
| **STAR Illumina** | 0.9842 ± 1.7E-3 | 0.9932 ± 1.0E-3 | 0.5 ± 0.0 |
| **STAR cutoff** | 0.99207 ± 3.7E-4 | 0.99784 ± 1.9E-4 | 0.5 ± 0.0 |
| **HISAT2 unfiltered** | 0.981 ± 0.012 | 0.985 ± 0.012 | 0.5 ± 0.0 |

JCC could retrieve true splice junctions not detected in the 50M reads data (Supplementary Figure S4). JCC’s AUPRC is 0.85 for the unfiltered STAR gold standard, but decreases when compared to the more strictly filtered gold standards (Table 3, “Real-world use case”). However, the performance of JCC in general is comparable to the performance of the No-Skill baseline classifier due to the high proportion of the positive class in the test data set.

### Scenario 3: Predicting hard-to-find junctions

#### Hypothetical use case

Hard-to-find junctions are defined as being 1) detected in 500M reads data, 2) not detected in 50M reads data, and 3) not annotated in the reference genome. These splice junctions represent rare or lower-expressed splicing events that are easily missed in datasets with lower sequencing depth. Similar to Scenario 2, we used the actual samples with low sequencing depth as JCC input (“Real-world use case”) and all splice junctions of interest as Splice AI and DeepSplice input (“Hypothetical use case”).

Both SpliceAI and DeepSplice (Table 4, “Hypothetical use case”) performed with a high AUPRC. SpliceAI outperformed DeepSplice with an AUPRC of 0.975 compared to 0.955 on the gold standard based on the unfiltered STAR results. Performance slightly improved with Illumina filtering (SpliceAI 0.978, DeepSplice 0.959). Due to the strictness of the STAR cutoff gold standard filtering, true splice junctions with high prediction scores might be filtered out. This can explain the overall worse performance on this gold standard (SpliceAI 0.89, DeepSplice 0.79). For the gold standard based on the HISAT2 results, both SpliceAI and DeepSplice outperformed the No-Skill baseline classifier (AUPRC 0.5) with an AUPRC of around 0.87.

**Table 4:** The mean area under the precision and recall curve (AUPRC) values ± 1 standard deviation per tool are shown for Scenario 3 “Predicting hard-to-find junctions” and gold standards STAR without filtering, STAR with Illumina filtering, STAR with cutoff > 10 uniquely aligned reads and HISAT2 without filtering averaged across all 4 × 10 subsampled 50M reads data. “No-Skill” describes a baseline classifier that predicts all splice junctions as positives.

|  | DeepSplice | SpliceAI | baseline classifier (“No-Skill”) |
| --- | --- | --- | --- |
| **STAR unfiltered** | 0.9546 ± 3.3E-3 | 0.9746 ± 1.9E-3 | 0.5 ± 0.0 |
| **STAR Illumina** | 0.9589 ± 3.5E-3 | 0.9781 ± 2.1E-3 | 0.5 ± 0.0 |
| **STAR cutoff** | 0.786 ± 0.012 | 0.893 ± 5.3E-3 | 0.5 ± 0.0 |
| **HISAT2 unfiltered** | 0.868 ± 0.09 | 0.878 ± 0.091 | 0.5 ± 0.0 |

For JCC performance, the AUPRC score would be highly misleading due to no true-positives and no false-positives at threshold 1, and therefore, precision being undefined. While JCC extracts some additional splice junctions not provided in the 50M reads input data, none of them are found in the STAR unfiltered gold standard. In Figure 2, we see that the mean of predictions for positives is lower than for negatives. Furthermore, the F1 scores of 0 for predictions thresholded at 0.5 reflect the poor predictive performance. In Supplementary Table S4 we report F1 scores across a range of thresholds from 0 to 1.

**Figure 2:**
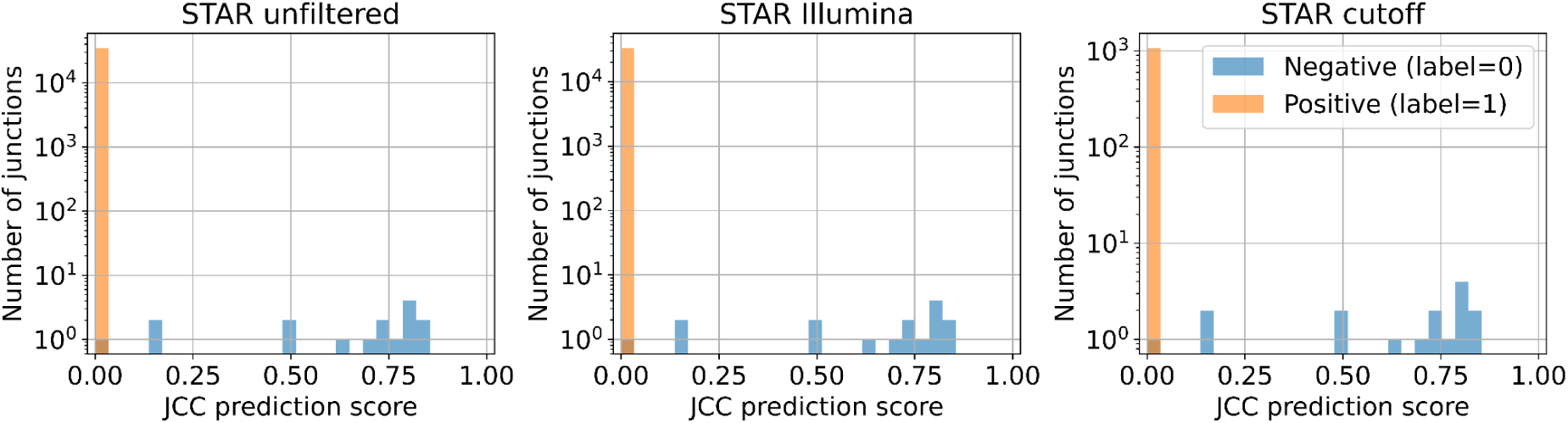
Distribution of JCC prediction scores for Scenario 3 “Predicting hard-to-find junctions” in Real-world use case according to STAR unfiltered, STAR Illumina filtered, and STAR cutoff filtered gold standards. The score distribution of splice junctions labeled as negatives (shown in blue) is compared to the score distribution of splice junctions labeled as positives (shown in orange). This is shown on sample J26675-L1_S1 subsample 0 as an example, but is similar for the other three samples and their subsamples.

Looking at the distribution of JCC prediction scores (Supplementary Figure S5), we find that splice junctions that are not annotated in the reference genome tend to have lower prediction scores than splice junctions that are not detected in the 50M reads data. However, SpliceAI and DeepSplice score distributions look similar for non-annotated splice junctions and splice junctions unsupported by 50M reads data (see Supplementary Text S1 “Tool score distributions”).

For a given tool, different types of splice junctions can be hard-to-find, depending on the input data. A tool, like JCC, relies on reference genome annotations to retrieve additional splice junctions. Our results highlight the limitations of both lower sequencing depth RNA-seq and reference genome annotations as input.

#### General recommendations

##### Which splice-aware aligner to use?

In our work, we used HISAT2 and STAR, as they are among the top splice-aware alignment tools. During the alignment process, both STAR and HISAT2 detected junctions from 50M reads data that were not detected from 500M reads data. This is due to technical noise introduced by these tools. While HISAT2 can be sensitive to the order of reads in the fastq file [41], STAR heavily penalizes motifs that differ from the canonical GT-AG splice junction. We suggest using STAR aligner over HISAT2, as HISAT2 introduces more technical noise (on average, 359 ± 21 noisy junctions in STAR vs. 7986 ± 1019 noisy junctions in HISAT2)

##### Should splice junctions be filtered using strict criteria?

Too strict filtering of splice junction candidates before running the splice junction prediction tools can result in the loss of potentially biologically valid low-confidence junctions. Therefore, we do not recommend any prior filtering, but instead hybrid filtering using the Illumina criteria combined with the splice prediction tool scoring.

##### Which splice prediction tool to use?

SpliceAI consistently outperforms DeepSplice and JCC across most evaluation scenarios. It is the recommended choice when high precision is required for filtering out noisy splice junctions. JCC is unique in its ability to rediscover previously unknown splice junctions by redistributing reads. However, it is performing poorly in retrieving unannotated splice junctions. The authors state that incomplete or wrong annotation or unreliable transcript estimates caused by sequence similarity or low read coverage of regions can lead to unreliable JCC predictions [32]. Furthermore, JCC requires both junction annotations and gene-spanning reads to be available; otherwise, predictions need to be imputed.

##### In which cases should one use a deep learning splice prediction tool?

For building the initial hypothesis, publicly available data sets with low sequencing depth can provide an easy and affordable entry point for alternative splicing detection. Also, as genome annotations are far from perfect and therefore often unable to capture the full diversity of splicing, sequence-based deep learning tools’ predictions combined with called experimental splice junctions from alignment tools could be used for improvement. Deep learning tools such as SpliceAI and DeepSplice can be used further for the following tasks. First, Scenario 1 suggests that they can be employed as an additional step in the standard RNA-seq pipeline to score and sensibly filter the spurious junctions that are still supported by RNA-seq reads. Second, Scenario 3 suggests that they can confirm a candidate, unannotated junction to be a real splice junction. However, a user should keep in mind that these deep learning models have been trained on DNA sequences and can not be used to reliably validate the usage of splice junctions for specific tissues, developmental stages, or environmental conditions. Thus, for comprehensive alternative splicing detection, higher sequencing depth data or complementary experimental approaches are still necessary. The experiment should be designed with sequencing depth of 150 to 200 million reads for low-expressed genes and 100 to 150 million reads for high-expressed genes [13], or it should be based on long-read sequencing like PacBio or Nanopore.

## Conclusion

We evaluated two approaches for alternative splicing event prediction from low sequencing depth data: starting from RNA-seq data (JCC) or genomic sequence (SpliceAI and DeepSplice). For this, we designed three distinct scenarios to investigate whether current tools can answer the three research questions.

We showed that all tools are bad at detecting noisy splice junctions. While SpliceAI mostly outperformed the other tools, the performance was influenced by the chosen aligner and filtering strategy, with HISAT2 introducing more spurious junctions (∼5%) than STAR (∼0.1%).

Furthermore, we showed that we can retrieve new splice junctions not detected in the 50M reads data, but with considerable limitations. SpliceAI and DeepSplice are not suitable for *de novo* splice junction detection, as they require junction coordinates to be provided by external annotations. Given these junction coordinates, both tools perform well, with SpliceAI outperforming DeepSplice. JCC, on the other hand, is designed for this task and attempts to recover novel splice junctions using deviations between predicted and observed coverage profiles. Its performance is limited in scenarios with low-depth sequencing data, and it only outperformed the No-Skill baseline classifier compared to the most stringent filtered STAR gold standard.

Finally, we showed that we can retrieve hard-to-find splice junctions, but only to a limited extent. These splice junctions, which are neither detected in samples with low sequencing depth nor annotated in the reference genome, represent rare or lowly expressed splicing events. While JCC could retrieve novel junctions, none were confirmed to be correct.

SpliceAI and DeepSplice substantially outperformed JCC, though in real-world scenarios, these models are limited by their dependency on pre-defined candidate splice junctions as input.

### Limitations

The study design has certain methodological limitations. First, the artificial decoy dataset might contain splice junctions. We mitigated this problem by filtering the generated GT-AG pairs for junctions that are neither annotated nor have ever been observed in any of the bigger public RNA-seq datasets (e.g., GTEx). Furthermore, though a huge fraction of annotated splice junctions are indeed GT-AG pairs, most randomly sampled GT-AG pairs might lack the surrounding ‘splicing code’ for proper recognition by the splicing machinery. Deep learning tools might easily detect the randomly sampled decoy splice junctions due to differing patterns in the surrounding sequence.

As mentioned in Methods, for the evaluation of whether the tools can predict true junctions without support in low sequencing depth data, DeepSplice and SpliceAI could not be run without information on positions from the high sequencing depth data, making it infeasible to run those tools for a user in a real-world use case.

### Future developments

Both RNA-seq data and genomic sequence contain information about the manifestation of alternative splicing events; however, tools that use both for splice junction prediction are still limited. TrueSight [42] was the first tool to use mapping quality from RNA-seq data and coding potential from reference genome sequences to detect splice junctions using logistic regression. However, the tool was published more than ten years ago and is currently not available. DARTS [43], while not a splice junction prediction tool but rather a differential splicing prediction tool, uses a deep neural network trained on manually compiled cis sequence features and mRNA levels of known trans RNA binding proteins. A Bayesian hypothesis testing statistical model integrates this prior probability prediction with empirical evidence from a specific RNA-seq dataset. However, manual feature selection, in opposition to learning features from the data, leads to bias and prevents the usage of novel biological features that are not yet well-studied. Thus, there is still a need for a deep learning tool that integrates information from both sequence and RNA-seq data to predict splicing using independently extracted features.

The next step should be the prediction of tissue- /condition-specific splicing. There are several developments in this direction. AbSplice predicts tissue-specific aberrant splicing from DNA sequence as well as RNA-seq data [44]. It combines scores from MMSplice and SpliceAI based on DNA sequence features with a tissue-specific splice site map and direct measures of aberrant splicing in clinically accessible tissues based on GTEx RNA-seq data. Pangolin [45] adopts a similar architecture as SpliceAI [29] to predict splice site usage across different tissues. Recently, the multi-transformer model TrASPr [23] was shown to outperform SpliceAI [29], Pangolin [45], and SpliceTransformer [46] in predicting tissue-specific PSI and dPSI from genomic sequences. However, the integration of RNA-seq and DNA sequence data for tissue- and condition-specific prediction of splicing and splice junction usage is still missing.

Besides the development of computational tools, sequencing technologies are also moving ahead. Further optimization of RNA-seq library preparation protocols and refinements in sequencing and mapping performance will facilitate improved splice junction prediction due to lower levels of technical noise. Given the falling sequ ncing costs, the more widespread availability of RNA-seq data from different organisms, tissues, and diseases will contribute to an increased understanding of the cell and tissue specificity of certain splice junctions.

Overall, while deep learning tools show promise in learning complex splicing rules for splice junctions from DNA sequence data, they can retrieve far fewer true splice junctions compared to junctions extracted from high sequencing depth data. However, we recommend sequence-based deep learning models like SpliceAI for initial hypothesis development and for supporting the splice-aware RNA-seq aligner as an additional filtering step in the standard RNA-seq pipeline, especially when sequencing depth is limited. For final confirmation of findings, though, further validation using higher sequencing depth data or complementary experimental approaches is necessary.

## Supporting information

Supplementary Table S4 - F1 scores at threshold

## Acknowledgements

The paper has also benefited greatly from my extensive discussions with Dr. Zoe Chervontseva, who made many suggestions.

## Ethics

The investigation conforms to the principles of the Declaration of Helsinki and the protocol was approved by the Ethics Committee of the University of Greifswald, Germany.

## Data and code availability

We used junctions from recount3 [38], adipose tissue data set [12] with GEO ID GSE46323, and a dilated cardiomyopathy data set, which is available upon request from University Medicine Greifswald. The code for tool evaluation and plots can be found at https://github.com/lenamariahackl/deep-sequencing-vs-DL. Junctions extracted by the aligners STAR and HISAT2 from 50M and 500M reads DCM data, as well as the prediction scores, are available at https://doi.org/10.5281/zenodo.16843445 [47]. The original RNA-seq data cannot be shared publicly for the privacy of individuals that participated in the study.

The data will be shared on reasonable request to the corresponding author.

