## Supplementary Files for "Detection of alternative splicing: deep sequencing or deep learning?"

### Supplement

#### Supplementary Text S1: Tool score distributions

Choice of strategy to combine donor and acceptor prediction scores into one score per splice junction is not highly important for SpliceAI, as only for one junction from the STAR 500M reads sample J26675-L1_S1 the predicted score difference between taking the mean and taking the maximum is more than 0.01. For JCC, the difference is only slightly more pronounced, with around 1% of predictions (3.469/289.226) differing more than 0.01, as shown in Supplementary Figure 6. For our evaluation, we use the maximum due to better performance.

The distributions of the resulting predictions differ. While scores range from 0 to 1 for tools SpliceAI and JCC, DeepSplice can return scores up to 1.7, as shown in Supplementary Figure 7. While the DeepSplice prediction values are approaching a Gaussian distribution, the distribution of JCC predictions is shifted towards the right, and the SpliceAI predictions appear to be beta-distributed with $\alpha = \beta = 0.5$.

#### Supplementary Text S2: Fitting the alpine bias model to run JCC

We set minLength = 8, maxLength = 21470 as the minimum and maximum length of coding single-isoform genes, and minCount = 0, maxCount = 88305 as the minimum and maximum read coverage of coding single-isoform genes. We kept nbrSubsample = 30 as described in the original paper. Parameters minsize and maxsize were chosen for the interval between them to contain at least the central 95 % of the fragment length distribution across samples (minsize=103, maxsize=16619).

#### Supplementary Tables:

| *Publication* | *Year* | *Reference* |
| --- | --- | --- |
| DNA-Level Splice Junction Prediction using Deep Recurrent Neural Networks | 2015 | [1] |
| "iSS-Hyb-mRMR": Identification of splicing sites using hybrid space of pseudo trinucleotide and pseudo tetranucleotide composition | 2016 | [2] |
| A computational approach for prediction of donor splice sites with improved accuracy | 2016 | [3] |
| Deep Learning Models Based on Distributed Feature Representations for Alternative Splicing Prediction | 2018 | [4] |
| **Discerning novel splice junctions derived from RNA-seq alignment: a deep learning approach** | 2018 | [5] |
| Human Splice-Site Prediction with Deep Neural Networks | 2018 | [6] |
| SpliceRover: interpretable convolutional neural networks for improved splice site prediction | 2018 | [7] |
| COSSMO: predicting competitive alternative splice site selection using deep learning | 2018 | [8] |
| SpliceVec: Distributed feature representations for splice junction prediction | 2018 | [9] |
| DeepSS: Exploring Splice Site Motif Through Convolutional Neural Network Directly From DNA Sequence | 2018 | [10] |
| Deep Splicing Code: Classifying Alternative Splicing Events Using Deep Learning | 2019 | [11] |
| DeepDSSR: Deep Learning Structure for Human Donor Splice Sites Recognition | 2019 | [12] |
| iSS-CNN: Identifying splicing sites using convolution neural network | 2019 | [13] |
| **Predicting Splicing from Primary Sequence with Deep Learning** | 2019 | [14] |
| Using the Chou's 5-steps rule to predict splice junctions with interpretable bidirectional long short-term memory networks | 2020 | [15] |
| EDeepSSP: Explainable deep neural networks for exact splice sites prediction | 2020 | [16] |
| Splice sites detection using chaos game representation and neural network | 2020 | [17] |
| Splice2Deep: An ensemble of deep convolutional neural networks for improved splice site prediction in genomic DNA | 2020 | [18] |
| SpliceFinder: ab initio prediction of splice sites using convolutional neural network | 2020 | [19] |
| Splice Junction Identification using Long Short-Term Memory Neural Networks | 2021 | [20] |
| DASSI: differential architecture search for splice identification from DNA sequences | 2021 | [21] |
| Spliceator: multi-species splice site prediction using convolutional neural networks | 2021 | [22] |
| SpliceViNCI: Visualizing the splicing of non-canonical introns through recurrent neural network | 2021 | [23] |
| Predicting RNA splicing from DNA sequence using Pangolin | 2022 | [24] |
| Splice-site identification for exon prediction using bidirectional LSTM-RNN approach | 2022 | [25] |
| DeepASmRNA: Reference-free prediction of alternative splicing events with a scalable and interpretable deep learning model | 2022 | [26] |
| Deep Splicer: A CNN Model for Splice Site Prediction in Genetic Sequences | 2022 | [27] |

| | | |
|---|---|---|
| EnsembleSplice: ensemble deep learning model for splice site prediction | 2022 | [28] |
| CNNSplice: Robust models for splice site prediction using convolutional neural networks | 2023 | [29] |
| DeepSplicer: An Improved Method of Splice Sites Prediction using Deep Learning | 2023 | [30] |

*Supplementary Table S1: List of publications introducing deep learning tools able to call splice junctions from sequence data input. Tools in highlighted publications were run in the course of this work.*

| **Term** | **Definition** |
|---|---|
| noisy junctions | umbrella term for both decoy splice junctions (investigated in Scenario 1a) with GT and AG motifs, but not observed in nature (see Section Decoy data set construction), and spurious junctions detected only in 50M reads data and not in 500M reads data (investigated in Scenario 1b). |
| spurious junctions | can be divided into technical noisy junctions found only in 50M reads data and not in filtered 500M reads data, and unsure junctions found in 50M reads data and unfiltered 500M reads data, but not in filtered 500M reads data. |
| hard-to-find junctions | splice junctions from the 500M reads gold standard that were not found in the subsampled 50M reads data and were not annotated in the reference genome. |

*Supplementary Table S2: Overview over most important definitions as used in this publication.*

Scenario 1: Detecting noisy junctions

a. Detecting junctions from a decoy dataset

| | | **STAR unfiltered** | **STAR Illumina** | **STAR cutoff** | **HISAT2 unfiltered** |
|---|---|---|---|---|---|
| **SpliceAI/ DeepSplice** | **positives** | 359 ± 21 | 3764 ± 60 | 60486 ± 1046 | 7986 ± 1019 |
| | **negatives** | 250835 ± 1940 | 247430 ± 1933 | 190708 ± 1244 | 178933 ± 2011 |

b. Detecting spurious junctions

| | | STAR unfiltered | STAR Illumina | STAR cutoff | HISAT2 unfiltered |
|---|---|---|---|---|---|
| **SpliceAI/ DeepSplice** | **positives** | 359 ± 21 | 3764 ± 60 | 60486 ± 1046 | 7986 ± 1019 |
| | **negatives** | 250835 ± 1940 | 247430 ± 1933 | 190708 ± 1244 | 178933 ± 2011 |
| **JCC** | **positives** | 359 ± 21 | 3764 ± 60 | 60486 ± 1046 | - |
| | **negatives** | 250835 ± 1940 | 247430 ± 1933 | 190708 ± 1244 | - |

Scenario 2: Predicting junctions that could be detected with higher sequencing depth

a. Real-world use case

| | | STAR unfiltered | STAR Illumina | STAR cutoff |
|---|---|---|---|---|
| **JCC** | **positives** | 448880 ± 12546 | 436090 ± 12381 | 204231 ± 1443 |
| | **negatives** | 60698 ± 1013 | 68945 ± 992 | 111120 ± 494 |

b. Hypothetical use case

| | | STAR unfiltered | STAR Illumina | STAR cutoff | HISAT2 unfiltered |
|---|---|---|---|---|---|
| **SpliceAI/ DeepSplice** | **positives** | 448880 ± 12546 | 436090 ± 12381 | 204231 ± 1443 | 249725 ± 9421 |
| | **negatives** | 448880 ± 12546 | 436090 ± 12381 | 204231 ± 1443 | 249725 ± 9421 |

Scenario 3: Predicting hard-to-find junctions

a. Real-world use case

| | | **STAR unfiltered** | **STAR Illumina** | **STAR cutoff** |
|---|---|---|---|---|
| **JCC** | **positives** | 39657 ± 3961 | 38117 ± 3917 | 1144 ± 81 |
| | **negatives** | 16 ± 0 | 16 ± 0 | 16 ± 0 |

b. Hypothetical use case

| | | **STAR unfiltered** | **STAR Illumina** | **STAR cutoff** | **HISAT2 unfiltered** |
|---|---|---|---|---|---|
| **SpliceAI/ DeepSplice** | **positives** | 39657 ± 3961 | 38117 ± 3917 | 1144 ± 81 | 9390 ± 2311 |
| | **negatives** | 39657 ± 3961 | 38117 ± 3917 | 1144 ± 81 | 9390 ± 2311 |

*Supplementary Table S3: Mean number of positive and negative splice junctions ± 1 standard deviation used for evaluation of the tools SpliceAI, DeepSplice and JCC for the three scenarios and different use cases averaged across all 4 × 10 subsampled 50M reads data.*

### Supplementary Figures:

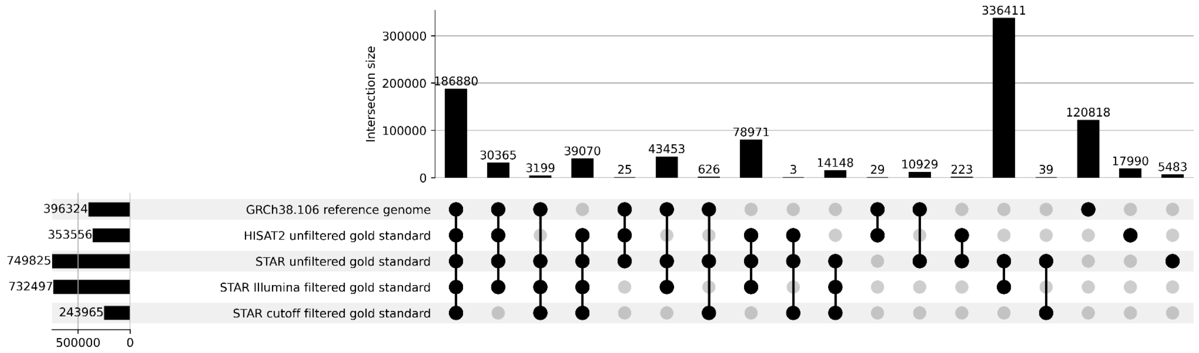


*Supplementary Figure S1: Upset plot comparing the GRCh38.106 reference genome splice* junction *annotations (reference) to the splice* junctions *called by HISAT2 on all four DCM RNA-seq samples with 500M reads (HISAT2 unfiltered gold standard), splice* junctions *called by STAR on those four DCM samples (STAR unfiltered gold standard), the STAR splice* junctions *filtered according to Illumina guidelines [31] (STAR Illumina filtered gold standard) and the STAR splice* junctions *filtered with a hard cutoff of at least 10 uniquely aligned reads (STAR cutoff filtered gold standard). This is shown for the combination of all 4 DCM samples.*

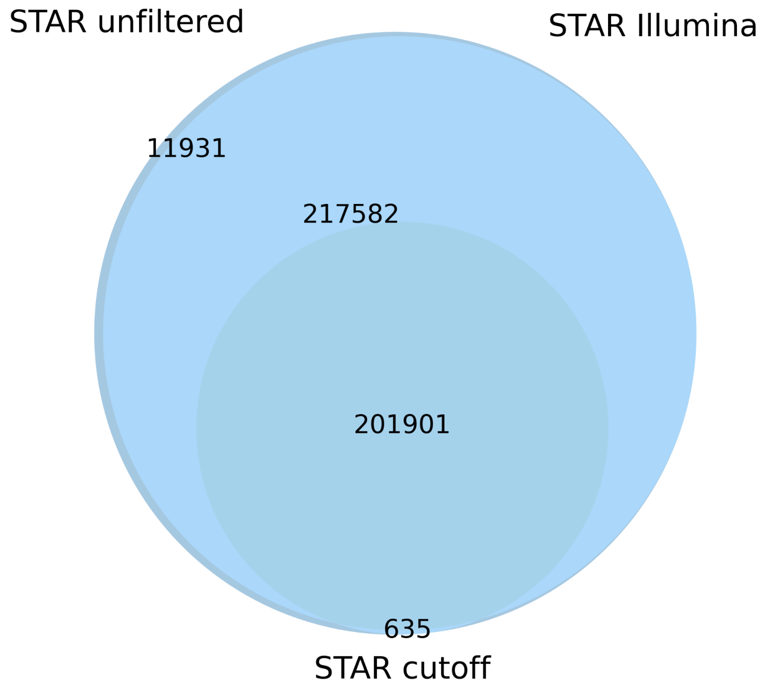


*Supplementary Figure S2: Venn diagram showing the overlap of the three different STAR junction gold standards after 1) no filtering, 2) filtering according to Illumina's suggested criteria, 3) keeping only junctions with more than cutoff 10 uniquely mapping reads. This is shown on sample J26675-L1_S1 as an example, but is similar for the other three samples.*

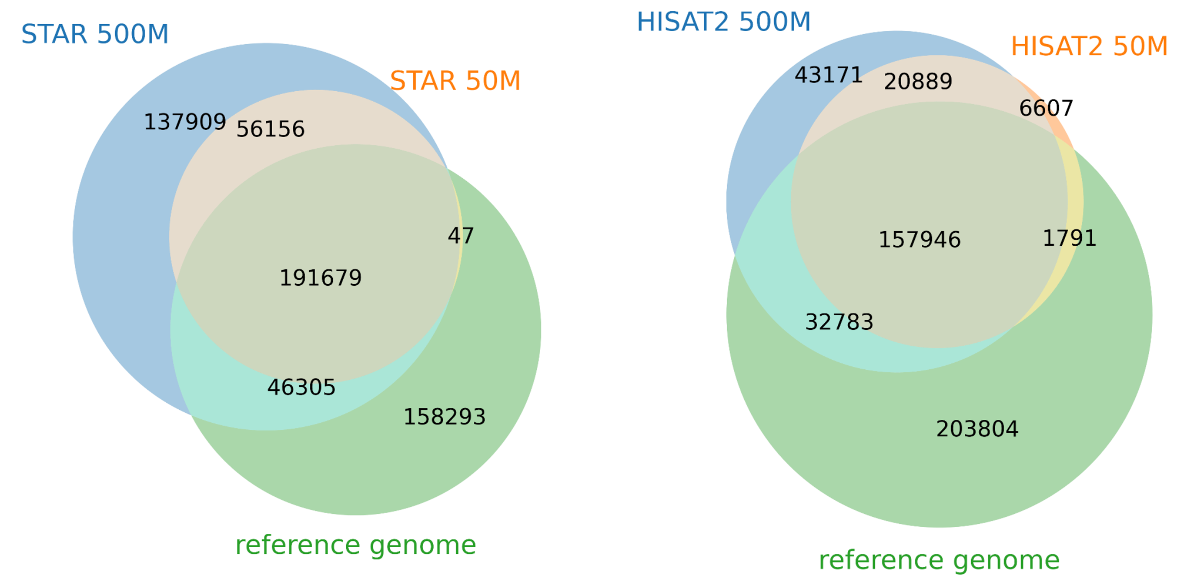


*Supplementary Figure S3: Venn diagrams showing the overlap of junctions from 500M reads gold standard aligned, junctions from randomly subsampled 50M reads data, and finally junctions from the GRCh38.106 reference genome. This is shown separately for aligners STAR and HISAT2 on sample J26675-L1_S1 subsample 0 as an example, but is similar for the other three samples and their subsamples.*

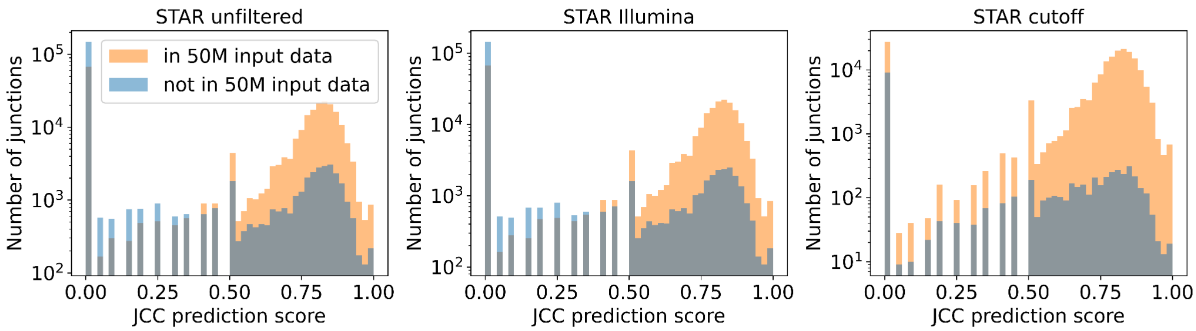


*Supplementary Figure S4: Distribution of JCC prediction scores for splice junctions labeled as positives according to STAR unfiltered, STAR Illumina filtered, and STAR cutoff filtered gold standards for Scenario 2 “Predicting junctions that could be detected with higher sequencing depth” in Real-world use case. The score distribution of splice junctions that were not contained in the 50M reads input data (shown in blue) is compared to the score distribution of splice junctions that were contained in the 50M reads input data (shown in orange). This is shown on sample J26675-L1_S1 subsample 0 as an example, but is similar for the other three samples and their subsamples.*

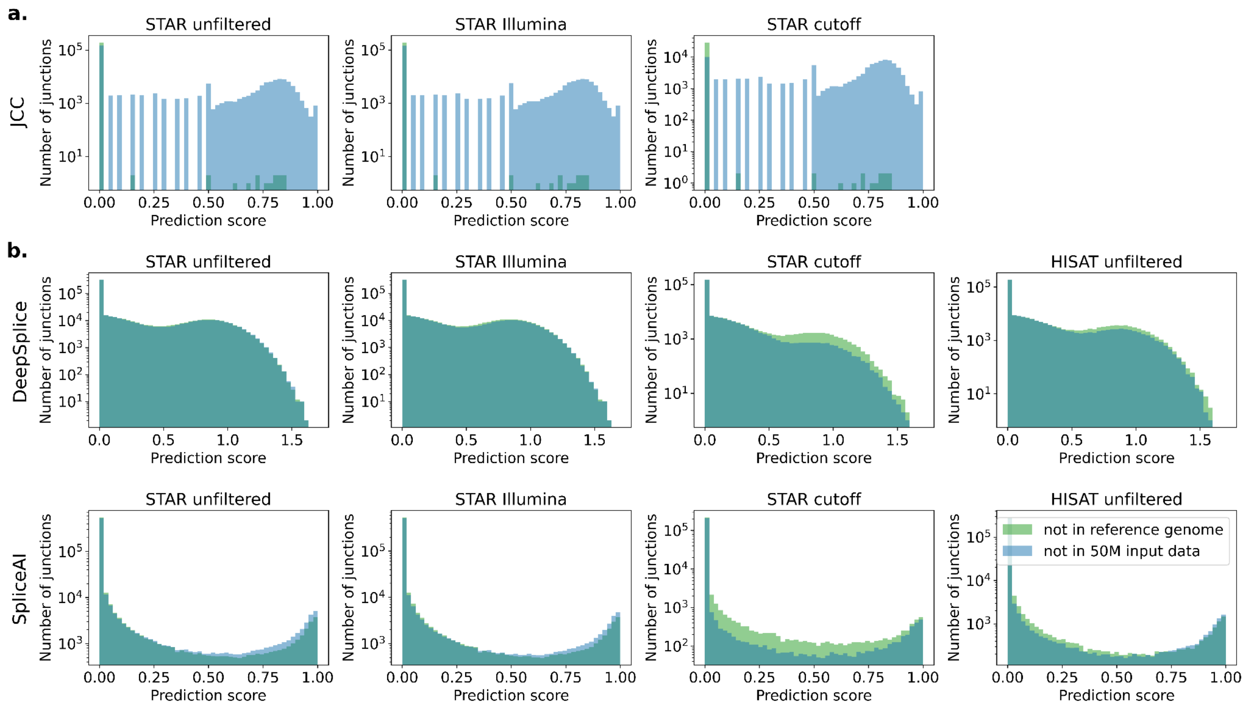


*Supplementary Figure S5: Distribution of prediction scores of JCC vs. DeepSplice vs. SpliceAI on gold standard data STAR unfiltered, STAR Illumina filtered, STAR cutoff filtered, HISAT2 unfiltered. The score distribution for splice junctions that were not found in the filtered 50M reads data (shown in blue) is compared to the score distribution for splice junctions that were not annotated in the reference genome (shown in green). In panel a., the prediction score distribution is visualized for Scenario 2: "Predicting junctions that could be detected with higher sequencing depth" Real-world use case, in panel b. for Scenario 2: "Predicting junctions that could be detected with higher sequencing depth" Hypothetical use case. This is shown on sample J26675-L1_S1 subsample 0 as an example, but is similar for the other three samples and their subsamples.*

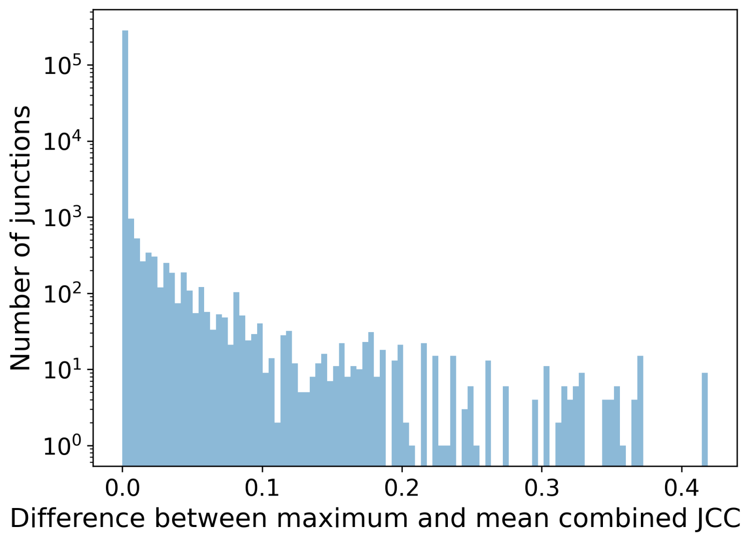


*Supplementary Figure S6: Distribution of the difference in JCC prediction scores between two strategies, maximum and mean, to combine donor and acceptor prediction scores into one score per splice junction for STAR 500M reads data. This is shown for sample J26675-L1_S1 as an example, but is similar for the other three samples.*

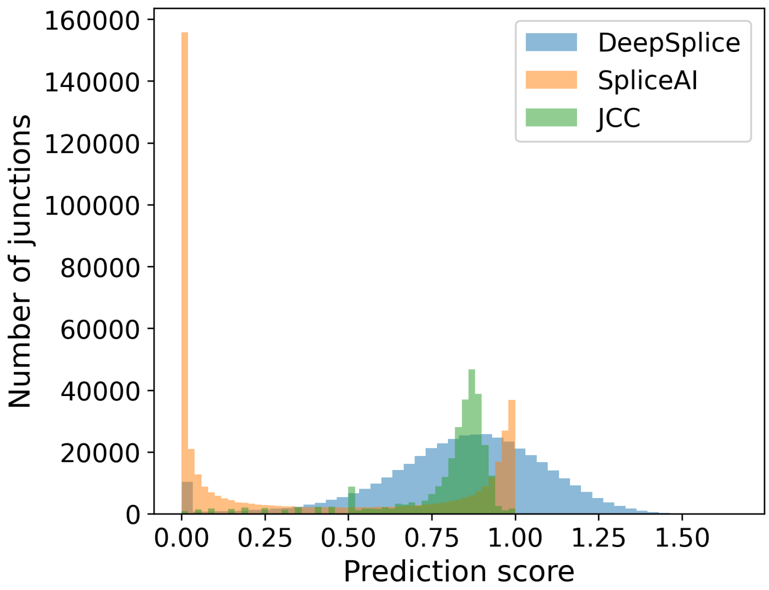


*Supplementary Figure S7: Distribution of the prediction scores of the three different tools DeepSplice, SpliceAI, and JCC, after running on the 500M reads RNA-seq data aligned with STAR. This is shown for sample J26675-L1_S1 as an example, but is similar for the other three samples.*
